# Multiple cis-regulatory elements collaborate to control *mdka* expression in telencephalic neural stem of adult zebrafish during constitutive and regenerative neurogenesis

**DOI:** 10.1101/2025.04.09.647643

**Authors:** Jincan Chen, Masanari Takamiya, Agnes Hendriks, Tanja Beil, Csilla Varnai, Nicolas Diotel, Sepand Rastegar

## Abstract

Zebrafish is a powerful animal model for studying nervous system regeneration due to its remarkable regenerative abilities and the availability of diverse molecular tools. After telencephalic brain injury, neural stem cells (NSCs) in the ventricular zone (VZ) become activated, proliferate, and generate new neurons essential for brain repair. However, the molecular mechanisms regulating these processes remain unclear. Here, we investigate the transcriptional regulation of *midkine-a* (*mdka*), a heparin-binding growth factor gene encoding the secreted protein Midkine-a (Mdka), which is upregulated after injury in radial glial cells (RGCs), the bona fide NSCs of the adult zebrafish telencephalon. Using genome-wide bioinformatic analysis, we identified six putative cis-regulatory elements (CREs) associated with *mdka*. Transgenic assays revealed that these CREs coordinate *mdka* expression during both development and regeneration. In the zebrafish embryo, CRE2, CRE3, CRE4, and CRE6 are required for EGFP expression in the nervous system, with CRE3 showing the strongest activity. In the adult telencephalon, CRE2, CRE4, and CRE6 are active in NSCs, with CRE2 best mimicking *mdka* expression at the ventricular zone. Importantly, individual CREs could not fully reproduce endogenous *mdka* expression, especially under regenerative conditions. In contrast, a combined CRE2346 construct closely recapitulated *mdka* expression in both the embryo and adult telencephalon under homeostatic conditions. These results suggest that *mdka* expression is controlled by a modular and cooperative cis-regulatory architecture that enables precise gene regulation during development, telencephalon homeostasis, and regeneration.

## Introduction

Understanding the molecular mechanisms underlying tissue and organ regeneration is critical for addressing the growing challenges posed by neurodegenerative diseases, such as dementia, Parkinson’s, and Alzheimer’s. These diseases place a significant medical and societal burden on an aging population, underscoring the urgent need for effective therapeutic strategies to address these conditions. Zebrafish has emerged as a key vertebrate model for studying regeneration due to its exceptional ability to regenerate tissues throughout both embryonic and adult stages [1–6]. Unlike most mammals, zebrafish can regenerate organs like the heart, fin, retina, spinal cord, and specific brain regions without scarring and obvious disabilities, which makes them invaluable for studying regenerative processes [2]. With over 70% genetic similarity to humans [7] and a vast number of mutants [8], transgenic lines [9–11], and genomic resources such as the DANIO-CODE Data Co-Ordination Center (DCC) (https://danio-code.zfin.org/), zebrafish provide a unique platform to study gene function, regulation, and cellular dynamics during embryonic development, adulthood under homeostatic and regenerative conditions [12–14]. In zebrafish, the adult telencephalon regenerates by activating dormant RGCs in the ventricular zone (VZ), which produce new neurons to replace those lost from injury [15, 16]. Recent transcriptomic studies have identified several genes significantly upregulated following injury [17–20], one of the most notable being *midkine-a* (*mdka*), a heparin-binding growth factor with multiple roles in development, repair and diseases [21–23]. Interestingly, *mdka* expression increases in quiescent RGCs after telencephalic injury, resembling the behaviour of *id1* (*inhibitor of differentiation 1*) [21], a transcriptional regulator downstream of the BMP signalling pathway [24, 25]. Both *mdka* and *id1* are predominantly expressed in quiescent type 1 cells (RGCs) and are absent in type 2 cells corresponding to actively dividing cells [21]. Both type 1 and type 2 cells express glial markers such as S100β and GFAP, while type 2 cells in addition express the proliferation marker PCNA (proliferating cell nuclear antigen). Lineage tracing experiments have demonstrated that type 2 cells differentiate into type 3 neuroblasts, which migrate and mature into specific neuronal types [26, 27].

Beyond the brain, *mdka* has been implicated in the regeneration of other tissues in zebrafish and other vertebrates, including the heart, fin, and retina, suggesting that it may play a common role in the regenerative process [28–35]. The zebrafish *midkine* family includes two genes: *mdka* and its paralog *midkine b* (*mdkb*), the result of a gene duplication event [36]. However, only *mdka* is significantly upregulated after injury, highlighting its specific role in regenerative responses [21]. This study investigates the transcriptional regulation of *mdka* in the zebrafish telencephalon, focusing on its regulation in NSCs during neurogenesis and regeneration. We aim to uncover the molecular mechanisms behind its transcription by identifying and characterizing cis-regulatory elements (CREs) that control *mdka* expression in both homeostatic and regenerative contexts.

Our findings, using genome-wide bioinformatic approaches and validation by transgenesis, reveal that multiple CREs work together to regulate *mdka* expression. In embryos, CRE2, CRE3, CRE4, and CRE6 are critical for nervous system expression, while in the adult telencephalon, CRE2, CRE4, and CRE6 are essential for expression in NSCs. Notably, none of the CREs alone were sufficient to fully recapitulate *mdka* expression in the embryonic nervous system or adult telencephalon. These findings highlight the complexity of *mdka* regulation, where individual CREs contribute differently to expression patterns depending on molecular and environmental signals. Furthermore, a transgenic EGFP reporter line containing CRE2, CRE3, CRE4, and CRE6 best mimicked endogenous *mdka* expression in both zebrafish embryos and the adult telencephalon under homeostatic and regenerative conditions, emphasizing the role of multiple molecular cues in *mdka* regulation.

## Results

### Identification and characterization of *mdka* CREs in the zebrafish embryo

To investigate the transcriptional regulation of *mdka*, we conducted a systematic search for CREs controlling *mdka* expression in zebrafish embryos at 24 hours post-fertilization (hpf) and in the ventricular zone of the adult telencephalon under both physiological and injury-induced conditions. Using data from the DANIO-CODE repository [14, 37], we identified six potential CREs located upstream, downstream, or in intergenic regions of the *mdka* coding sequence, as shown in **Figure 1**. These regions were selected based on chromatin accessibility, as assessed by ATAC-seq (Assay for Transposase-Accessible Chromatin with sequencing) across zebrafish developmental stages, from dome to long-pec. We also considered sequence conservation with the carp genome [38], indicated by blue boxes. Based on epigenetic features provided in the DANIO-CODE resource, including DNA methylation and histone modifications, the potential CREs were classified as promoters or enhancers and categorized by their activation state (repressed, quiescent, or activated) and strength (weak or strong). To validate their regulatory activity *in vivo*, each CRE was cloned upstream of a 1.0-kb *gata2* minimal promoter, corresponding to the most truncated basal promoter (P5) described by Meng et al. (1997) [39]. This promoter has been subsequently validated for enhancer assays by Navratilova et al. (2009) and Zhang et al. (2020) [25, 40], and was used here to drive EGFP expression (**Figure 2A**). These constructs were introduced into zebrafish germline cell to generate stable transgenic lines, *Tg(CREx-gata2aPR:EGFP)*, where CREx represents one of the identified CREs (**Figure 2A**). Five of the six CREs (CRE2-6) drove expression in 24 hpf zebrafish embryos, displaying partially overlapping patterns with endogenous *mdka* expression (**Figure 2B-G; B’-G’; Figure S1A, B**). Among them, *Tg(CRE3-gata2aPR:EGFP)* most closely recapitulated *mdka* expression (**Figure 2D, D’** and **Table 1**). In this line, EGFP was detected along the spinal cord, tail/fin fold, and brain (**Figure 2D, D’; Table 1; Figure S1A** shows a representative 24 hpf zebrafish embryo with specific anatomical regions). However, none of the individual CREs fully replicated the endogenous *mdka* expression pattern (**Figure 2; Table 1**). No EGFP expression was detected in the stable lines generated with CRE1 **(Figure 2B, B’)**. Several explanations could account for this, including the possibility that CRE1 functions as a silencer. This hypothesis will be briefly discussed in the relevant section. Additionally, each CRE was tested in combination with three different promoters: the *ctgf* promoter [41], the zebrafish synthetic promoter (zsp) [42], and the *mdka* intrinsic promoter (mdkaPR, this study). EGFP expression in these transient transgenics closely resembled that observed using the *gata2a* promoter (**Figure S2**), supporting the conclusion that EGFP expression is primarily driven by the CREs rather than by the specific promoter used. Given the consistent expression patterns observed with the *gata2a* promoter, all subsequent characterization of the CREs was carried out using the *gata2aPR:EGFP* construct.

**Figure 1.**
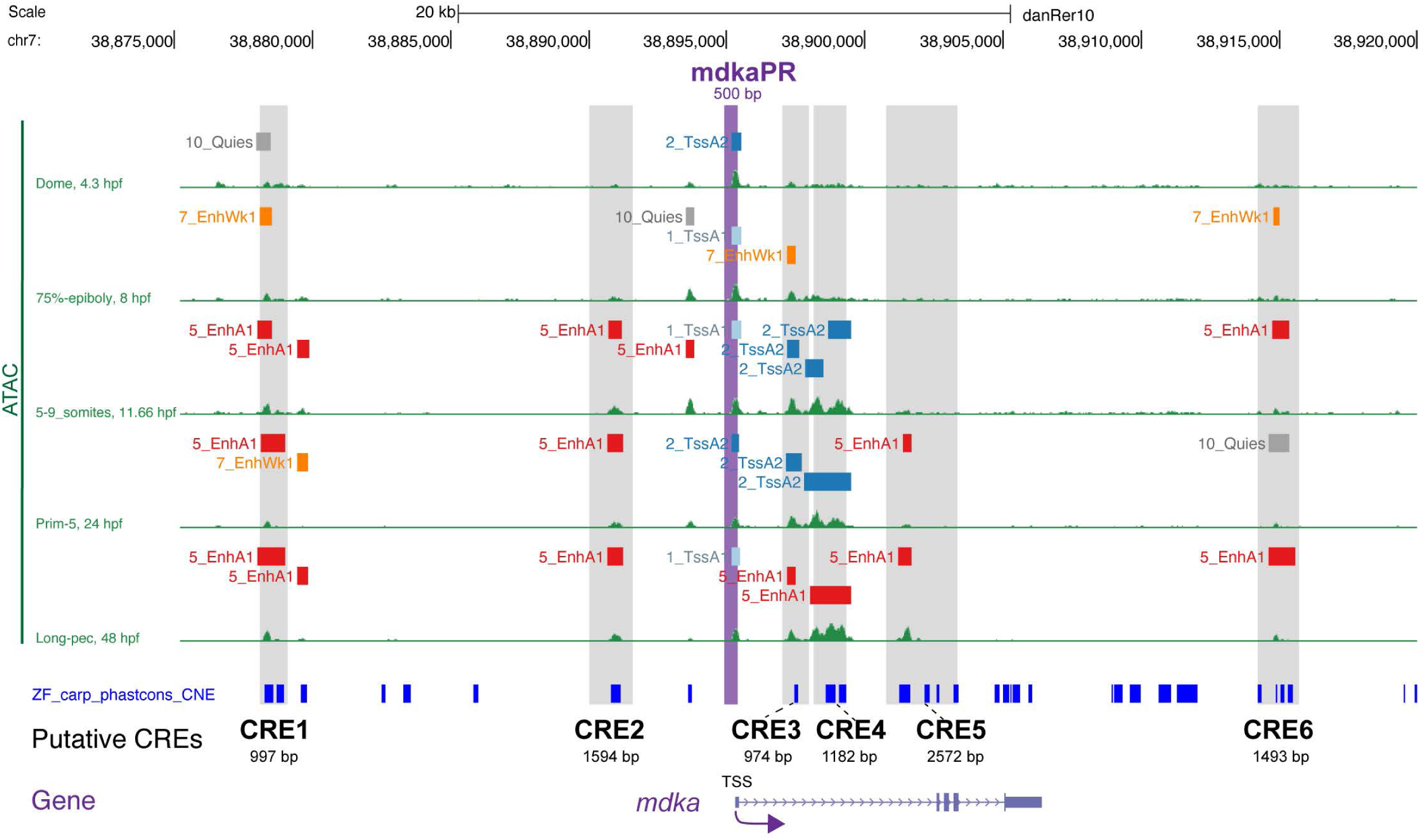
Identification of potential *mdka* regulatory sequences based on embryonic developmental ATAC-seq data. Snapshot of UCSC genome browser tracks displaying ATAC-seq data from different embryonic developmental stages (Dome, 75%-epiboly, 5-9 somites, Prime-5 and Long-pec) across a 45-kilobase region (chr7: 38,875,000-38,920,000; danRer10 assembly). Green peak profiles represent open chromatin regions identified by ATAC-seq. Color-coded horizontal bars indicate the genomic position of putative regulatory elements based on PADRES (Predicted ATAC-seq-supported developmental regulatory elements): promoters (1_TssA and 2_TssA2, blue), enhancers (5_EnhA1, red; 7_EnhWk1, orange), and other elements (e.g., 10_Quies, grey). The grey box highlights six putative *mdka* CREs (CRE1 to CRE6) identified in this study, while the magenta box marks the putative *mdka* promoter region. The relative positions of all CREs are shown in reference to the *mdka* gene and its transcription start site (TSS). PhastCons conservation scores derived from zebrafish-carp genome alignment highlights conserved non-coding regions (CNEs) associated with the *mdka* locus.

**Figure 2.**
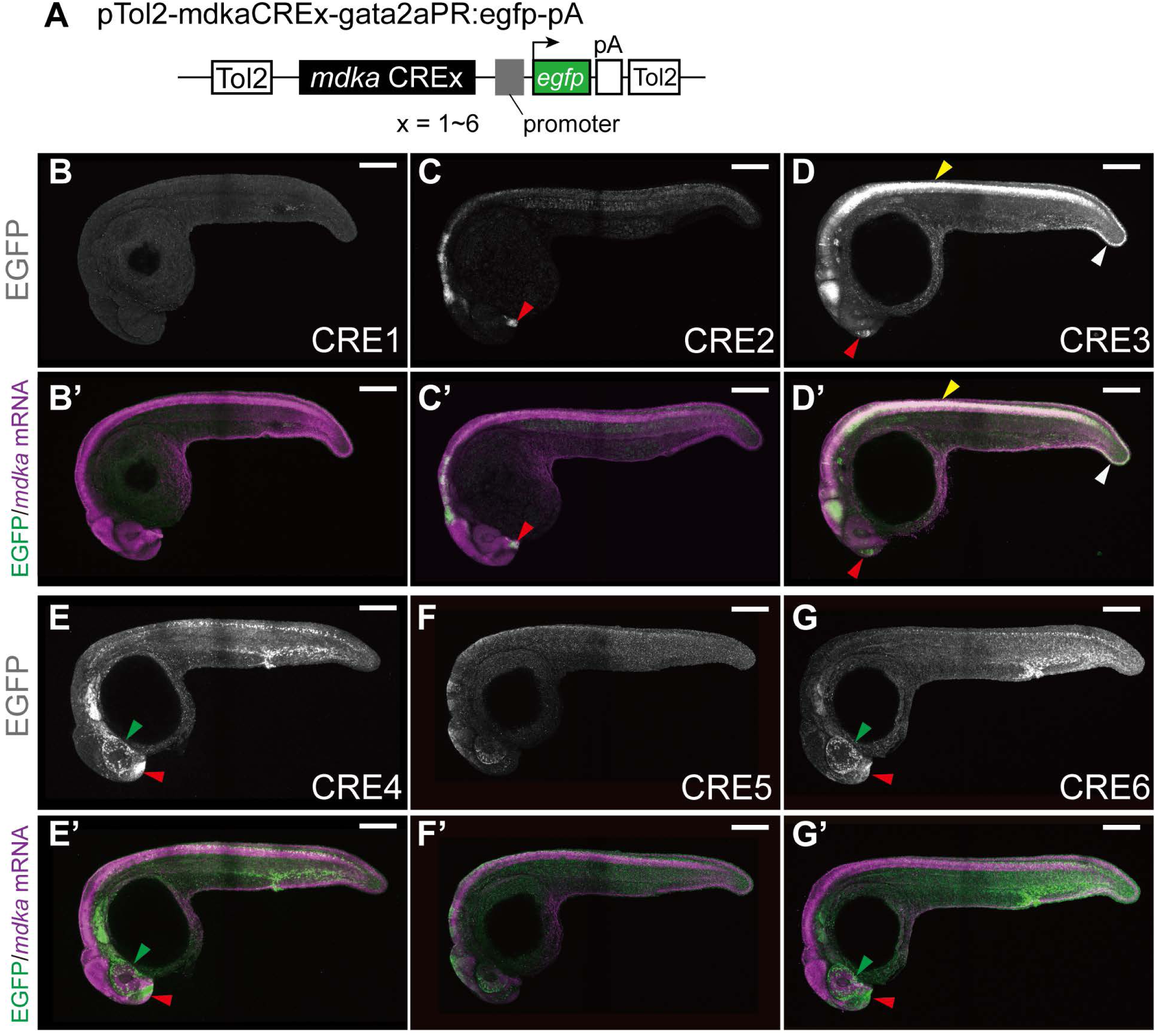
Stable EGFP transgenic line generated using potential *mdka* CREs. **A**, Schematic of the plasmid pTol2-mdkaCREx-gata2aPR:egfp-pA, where CREx represents one of the six putative CREs (CRE1 to CRE6). **B-G**, Stable EGFP reporter expression driven by *mdka* CRE1 (B), CRE2 (C), CRE3 (D), CRE4 (E), CRE5 (F), CRE6 (G) in embryo at 24 hours post-fertilization (hpf). **B’-G’**, Merged images showing *mdka* mRNA (magenta) and EGFP (green) expression in the corresponding CRE transgenic lines. *Tg(CRE3-gata2aPR:EGFP)* embryo exhibited strong co-localization of *mdka* mRNA and EGFP signals in the spinal cord (yellow arrowheads, D, D’) and fin fold region (white arrowheads, D, D’). *Tg(CRE4-gata2aPR:EGFP)* and *Tg(CRE6-gata2aPR:EGFP)* embryos showed strong activity in the retina (green arrowheads) and brain regions (red arrowheads, E, E’, G, G’), respectively. *Tg(CRE1-gata2aPR:EGFP)* showed no detectable EGFP expression (B, B’). Scale bar: 200 μm.

**Table 1.** Summary of EGFP expression patterns driven by individual CREs in 24 hpf embryos.

|  | Endogenous | CRE1 | CRE2 | CRE3 | CRE4 | CRE5 | CRE6 |
| --- | --- | --- | --- | --- | --- | --- | --- |
| Brain | ++ | - | + | + | + | + | + |
| Retina | ++ | - | + | ++ | ++ | + | ++ |
| Spinal cord | ++ | - | + | ++ | + | + | + |
| Notochord | + | - | + | - | - | - | - |
| Fin fold/Tail | ++ | - | - | ++ | + | - | + |
| Somite/Muscle | ++ | - | + | - | ++ | - | ++ |

Note: -, no detectable expression; +, weak expression; ++, strong expression.

### Characterization of CREs driving *mdka* expression in adult NSCs

In the adult zebrafish telencephalon, EGFP expression was detected in four out of six transgenic lines generated (**Figure 3C, C’; D, D’; E, E’; G, G’**). No expression was observed for CRE1 and CRE5 (**Figure 3B, B’; F, F’**). CRE3 drove strong ectopic EGFP expression in the dorsal pallium (Dp) and lateral pallium (Dl), whereas no expression was detected in the dorsomedial region (Dm) of the VZ (**Figure 3D, D’**). In contrast, CRE2, CRE4, and CRE6 directed EGFP expression in the VZ of the telencephalon, where RGCs reside (**Figure 3C, C’; E, E’; G, G’**, and **Table 2**). To determine if EGFP transgene expression accurately reflects endogenous *mdka* expression, we compared it with endogenous *mdka* expression at single-cell resolution (**Figure 3H-J**). In CRE2, CRE4 and CRE6 transgenic lines, EGFP-positive cells were present along the VZ with endogenous *mdka*-positive RGCs (**Figure 3H-J’’**). Quantification of confocal stack images showed that roughly half of *mdka*-positive RGCs co-expressed EGFP in each CRE transgenic lines, with no significant difference among the three CRE lines (**Figure 3K**). In the CRE6 transgenic line, EGFP expression was also observed outside of VZ, where endogenous *mdka* is not expressed, suggesting that CRE6 activity leads to ectopic transgene expression (**Figure 3J’’**, white arrows). Since we previously reported that *mdka* expression is predominant in quiescent RGCs compared to activated RGCs [21], we next exanimated whether CRE2, CRE4 and CRE6 drive EGFP expression in similar manner. To investigate this, we analyzed the expression of the RGC marker S100β and the proliferation marker PCNA in EGFP-positive cells of each CRE transgenic line (**Figure 3L**). All CRE transgenic lines exhibited robust EGFP expression, primarily in quiescent type 1 RGCs marked by S100β+/PCNA−, closely resembling the endogenous *mdka* expression (**Figure 3M; Figure S1C; Figure S3A-A’’’, B-B’’’**). Although each CRE captured specific aspects of endogenous *mdka* expression, none of them alone fully recapitulated its complete spatial pattern in the adult telencephalon. A similar discrepancy was observed during embryonic development at 24 hpf (**Figure 2**). While *Tg(CRE2-gata2aPR:EGFP)*, *Tg(CRE4-gata2aPR:EGFP)*, and *Tg(CRE6-gata2aPR:EGFP)* each reflect certain aspects of *mdka* expression, their activity was incomplete in both spatial distribution and intensity. Interestingly, CREs that were strongly active in embryonic central nervous system (**Figure 2C**, **C’**) also exhibited activity in the adult telencephalon (**Figure 3C, E and G**). However, their expression patterns differed from that of the endogenous *mdka* mRNA, suggesting that distinct regulatory elements control *mdka* expression at different developmental stages. These findings indicate that full *mdka* expression likely requires a combination of multiple CREs, and that these combinations may vary between embryonic and adult stages, particularly in the adult telencephalon.

**Figure 3.**
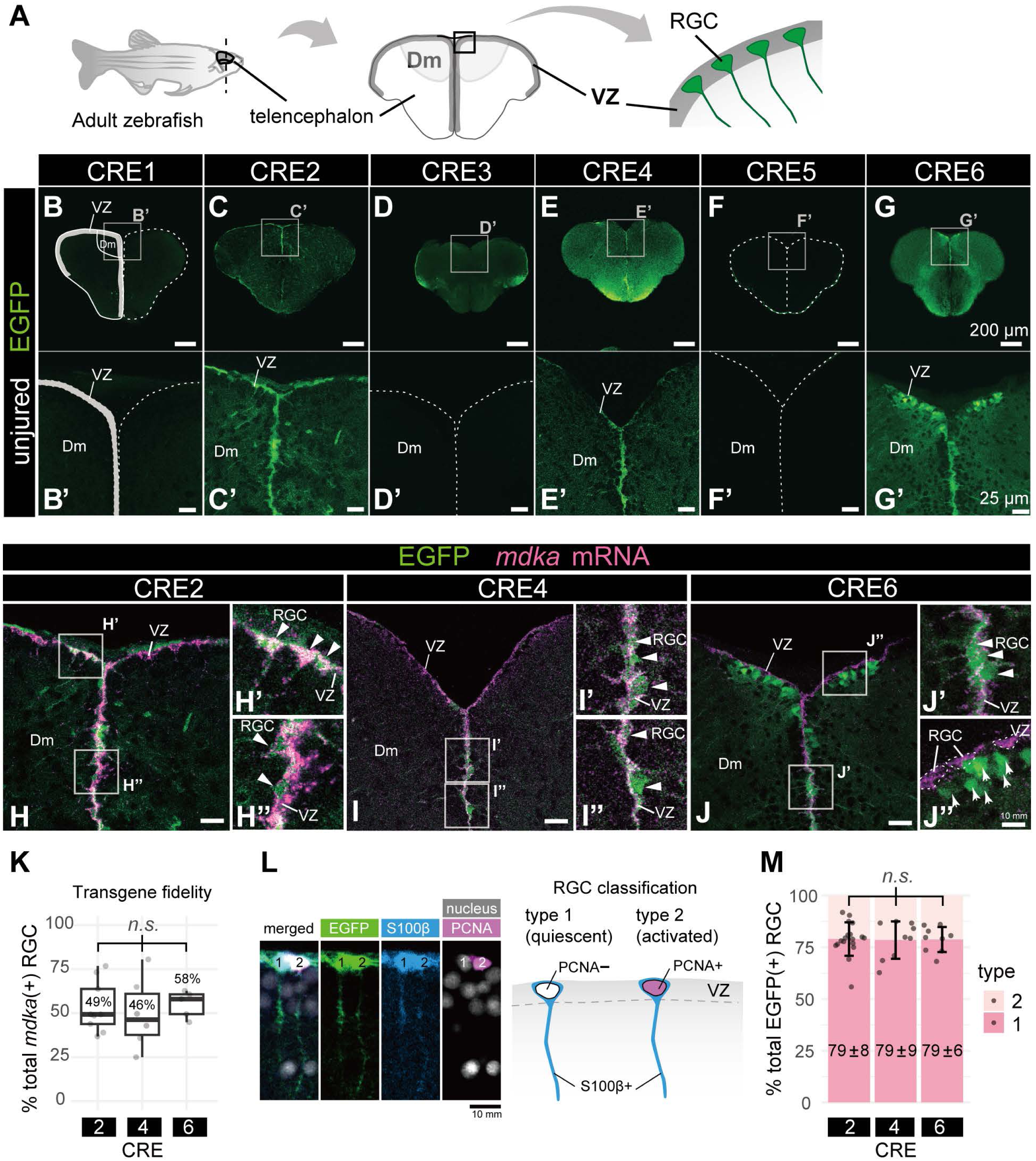
The *mdka* transgenic lines, CRE2, CRE4 and CRE6, drive radial glial EGFP expression in the adult telencephalon during constitutive neurogenesis. **A**, Schematic illustrating the orientation of transverse sections through the telencephalon, with reference to the location of radial glial cells (RGCs). Dm: dorsomedial pallium; VZ: ventricular zone; RGCs: radial glial cells. **B-G’**, EGFP reporter expression in the adult telencephalon across various *mdka* CRE transgenic lines: CRE1 (B-B’. *n* = 3 brains), CRE2 (C-C’, *n* = 4), CRE3 (D-D’, *n* = 6), CRE4 (E-E’, *n* = 3), CRE5 (F-F’, *n* = 4), and CRE6 (G-G’, *n* = 3). The location of VZ is indicated on the left hemisphere of CRE1 telencephalon, where EGFP expression is absent (B-B’). Rectangles in B-G indicate magnified regions shown in B’-G’, respectively. Dashed lines outline the telencephalon contour (B-G). **H-J’’,** Magnified transverse views of the Dm region showing EGFP (green) and *mdka* mRNA expression (magenta) in three transgenic lines with radial glial EGFP expression: *Tg(CRE2-gata2aPR:EGFP)*(H-H’’), *Tg(CRE4-gata2aPR:EGFP)*(I-I’’), and *Tg(CRE6-gata2aPR:EGFP)* (J-J’’). White arrowheads highlight RGCs of *mdka* mRNA and EGFP co-localization, reflecting faithful activity of CRE2, CRE4 and CRE6. Arrows in the panel J’’ indicate the ectopic EGFP expression of CRE6 located outside of VZ. **K**, The fidelity of CRE2, CRE4 and CRE6 activity was evaluated by quantifying the percentage of EGFP-positive cells among endogenous *mdka*-positive RGCs with the confocal images as shown the panels H-J. Results are shown as boxplots with the median values. No significant difference was observed between different CREs (*ANOVA* test, *p* = 0.84). **L**, S100β+ EGFP cells are classified either as type 1 (quiescent, PCNA-negative) or type 2 (activated, PCNA-positive) based on PCNA expression. **M**, Quantification of type 1 and type 2 RGC population in CRE2, CRE4, and CRE6. The mean ± standard deviation (error bar) of type 1 RGCs is expressed as a percentage of the total RGC population. No significant difference was observed between different CREs (*n.s.*; *Kruskal-Wallis* test, *p* = 0.97). Representative confocal images corresponding to this quantification are provided in Supplementary Figure S3A-A’’’, B-B’’’ and Figure S4A-A’’’, C-C’’’, showing co-immunostaining for EGFP, S100β, and PCNA. Each data point represents an individual brain slice. Scale bar: 200 μm (B-G), 25 μm (B’-G’, H-J) and 10 µm (H’-H’’, I’-I’’, J’-J’’ and L). *n* = 3 brains (CRE1; B, B’), *n* = 6 (CRE2; C, C’, H-H’’, K, M), *n* = 6 (CRE3; D, D’), *n* = 3 (CRE4; E, E’, I-I’’, K, M), *n* = 4 (CRE5; F, F’), *n* = 3 (CRE6; G, G’, J-J’, K, M).

**Table 2.** Summary of EGFP expression patterns driven by individual CREs in the adult telencephalon.

|  | Endogenous | CRE1 | CRE2 | CRE3 | CRE4 | CRE5 | CRE6 |
| --- | --- | --- | --- | --- | --- | --- | --- |
| Dm | ++ | - | ++ | - | + | - | + |
| DI | ++ | - | + | + | + | - | + |
| Dp | + | - | - | ++ | - | - | - |
| Vd | + | - | + | - | - | - | - |
| Vv | + | - | + | - | ++ | - | - |

Note: -, no detectable expression; +, weak expression; ++, strong expression. Abbreviations: Dm, dorsomedial pallium; Dp, dorsal pallium; DI, lateral pallium; VZ, ventricular zone; Vd, dorsal nucleus of ventral telencephalon; Vv, ventral nucleus of ventral telencephalon.

### Identification of injury-responsive CREs regulating *mdka* expression in the adult telencephalon

Since *mdka* expression is significantly upregulated during telencephalon regeneration following brain injury, particularly in type 1 RGCs [21], we subjected all six *mdka* CRE transgenic lines to telencephalon injury (**Figure 4B-G, B’-G’**). All injury-related analyses were performed at 5 days post-lesion (dpl), based on prior RNA-sequencing and RT-qPCR data showing that *mdka* expression peaks at this time point in the injured hemisphere, with a 1.37-fold increase compared to the uninjured side (*p* = 2.68 × 10⁻⁹) [21]. This time point therefore provided an optimal window to evaluate injury-induced regulatory activity of the CREs. The injury was induced in the right hemisphere using a 30G syringe needle inserted through the skull [27, 43], while the uninjured contralateral hemisphere served as an internal control (**Figure 4A**). Following injury, EGFP reporter expression remained absent in the VZ of the Dm region in transgenic lines carrying CRE1, CRE3, and CRE5 (**Figure 4B, B’, D, D’, F,** and **F’**), consistent with the lack of expression observed under homeostatic conditions (**Figure 3B, B’, D, D’, F,** and **F’**). In contrast, transgenic lines with CRE2 (**Figure 4C, C’**), CRE4 (**Figure 4E, E’**), and CRE6 (**Figure 4G, G’**) showed EGFP expression in the Dm of the VZ.

**Figure 4.**
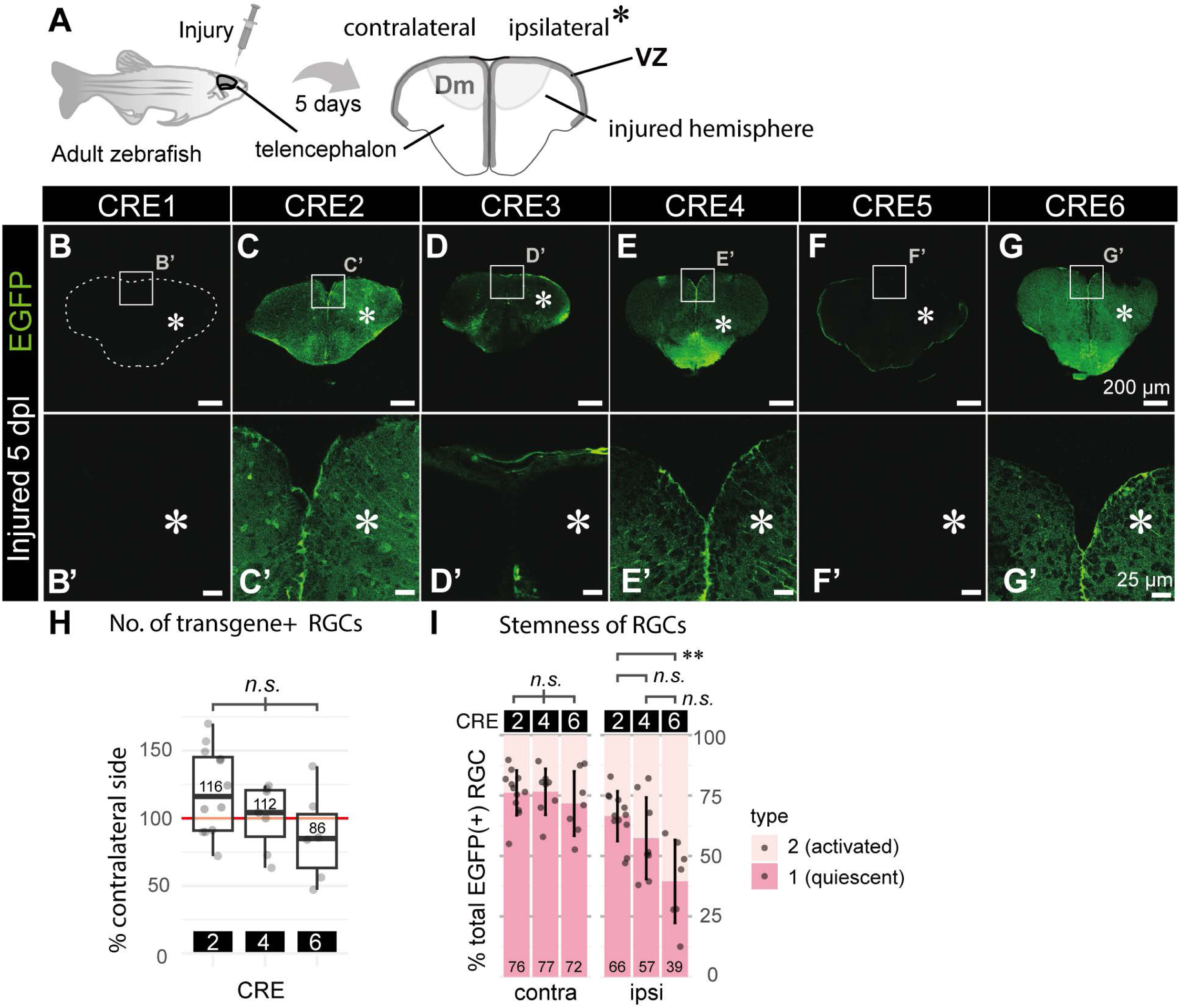
During regenerative neurogenesis, *mdka* CRE2, CRE4 and CRE6, maintain radial glial EGFP expression, though show variability in the injured brain hemisphere. **A**, Schematic illustrating the experimental design and orientation of telencephalic brain sections. Adult zebrafish were subjected to stab injury in the right hemisphere of the telencephalon, and brains were collected and fixed at 5 days post-lesion (dpl). **B-G’**, EGFP expression from stable transgenic lines for CRE1 (B, B’), CRE2 (C, C’), CRE3 (D, D’), CRE4 (E, E’), CRE5 (F, F’), and CRE6 (G, G’) is shown for both contralateral and ipsilateral sides (asterisks indicate ipsilateral/injured side) of the telencephalon. Rectangles in B-G indicate magnified regions shown in B’-G’, respectively. Dashed lines mark the telencephalon contour (B)**. H,** Quantification of EGFP-positive and S100β-positive RGCs. The number of EGFP-positive RGCs in the injured (ipsilateral) hemispheres of CRE2, CRE4, and CRE6 transgenic lines is shown as a percentage relative to the corresponding contralateral side in boxplots. No significant differences were observed among CREs (*ANOVA*, *p* > 0.05). **I,** Stemness of EGFP-positive RGCs in CRE2, CRE4, and CRE6 was assessed. Relative proportions of type 1 (quiescent, S100β⁺/PCNA⁻) and type 2 (activated, S100β⁺/PCNA⁺) RGCs in contralateral and ipsilateral hemispheres are presented as stacked bar plots. **While no statistical differences were found among CREs in the contralateral hemisphere, a significant difference was observed between CRE2 and CRE6 in the ipsilateral (injured) hemisphere (*p* < 0.01). Representative confocal images used for quantifications in panels H and I are provided in Supplementary Figures S3C-D’’’ and S4B-D’’’, showing co-immunostaining for EGFP, S100β, and PCNA. Each data point represents an individual brain slice. Scale bars: 200 μm (B-G); 25 μm (B’-G’). Sample sizes: *n* = 3 brains (B, B’), *n* = 4 (C, C’), *n* = 5 (D, D’), *n* = 3 (E, E’), *n* = 4 (F, F’), *n* = 3 (G, G’), *n* = 3 for each CRE (H, I).

However, for all three CREs, the number of EGFP-positive RGCs in the injured hemisphere remained largely unchanged compared to the uninjured contralateral side, with no significant differences between the CREs (**Figure 4H**). Assessment of RGC stemness, based on the proportion of type 1 RGCs among EGFP-positive cells, revealed differences between CREs in the injured (ipsilateral) brain hemisphere. Specifically, CRE2 showed a significantly higher proportion of quiescent type 1 RGCs compared to CRE6 (**Figure 4I**, *p* < 0.01). In contrast, on the contralateral (uninjured) side, approximately three-quarters of EGFP-positive RGCs remained quiescent across all CRE lines, consistent with the proportions observed under homeostatic conditions (**Figure 3M**; **Figure 4I**). These results suggest that none of the individual CREs contain a strongly injury-responsive regulatory element sufficient to upregulate *mdka* expression during regeneration. Similar to the regulation observed under physiological conditions, it is likely that multiple CREs must act in combination to mediate a full regenerative response.

### Cooperative CRE activity enhances faithful transgene expression in embryos, adult RGCs, and in response to injury

Analysis of individual stable transgenic lines showed that none of the six CREs alone fully recapitulated the spatial expression pattern of *mdka* throughout embryonic development and adulthood, under either homeostatic or regenerative conditions. To investigate potential cooperative interactions among these CREs, we generated a combined CRE2346 transgenic reporter construct (**Figure 5A**). This construct integrates CRE3, which showed strong regulatory activity during embryogenesis, together with CRE2, CRE4, and CRE6, which were effective in the adult telencephalon. Under homeostatic conditions, the CRE2346 construct drove EGFP expression patterns that closely mimicked endogenous *mdka* expression in embryos (**Figure 5B-B’’**). In the adult telencephalon (**Figure 5C, D-D’’**), CRE2346 mediated specific EGFP expression in the Dm of the VZ (white arrowheads, **Figure 5D’**), displaying RGC morphology and co-expression with the RGC marker S100β. Transgene fidelity assessment revealed that CRE2346 was active in the majority (median 88%) of *mdka*-positive RGCs, demonstrating significantly higher coverage than any individual CRE (**Figure 5E**). However, analysis of RGC stemness, assessed by immunolabeling for PCNA and S100β, showed no significant effect of the CRE combination (**Figure 5F-F’’’, G**).

**Figure 5.**
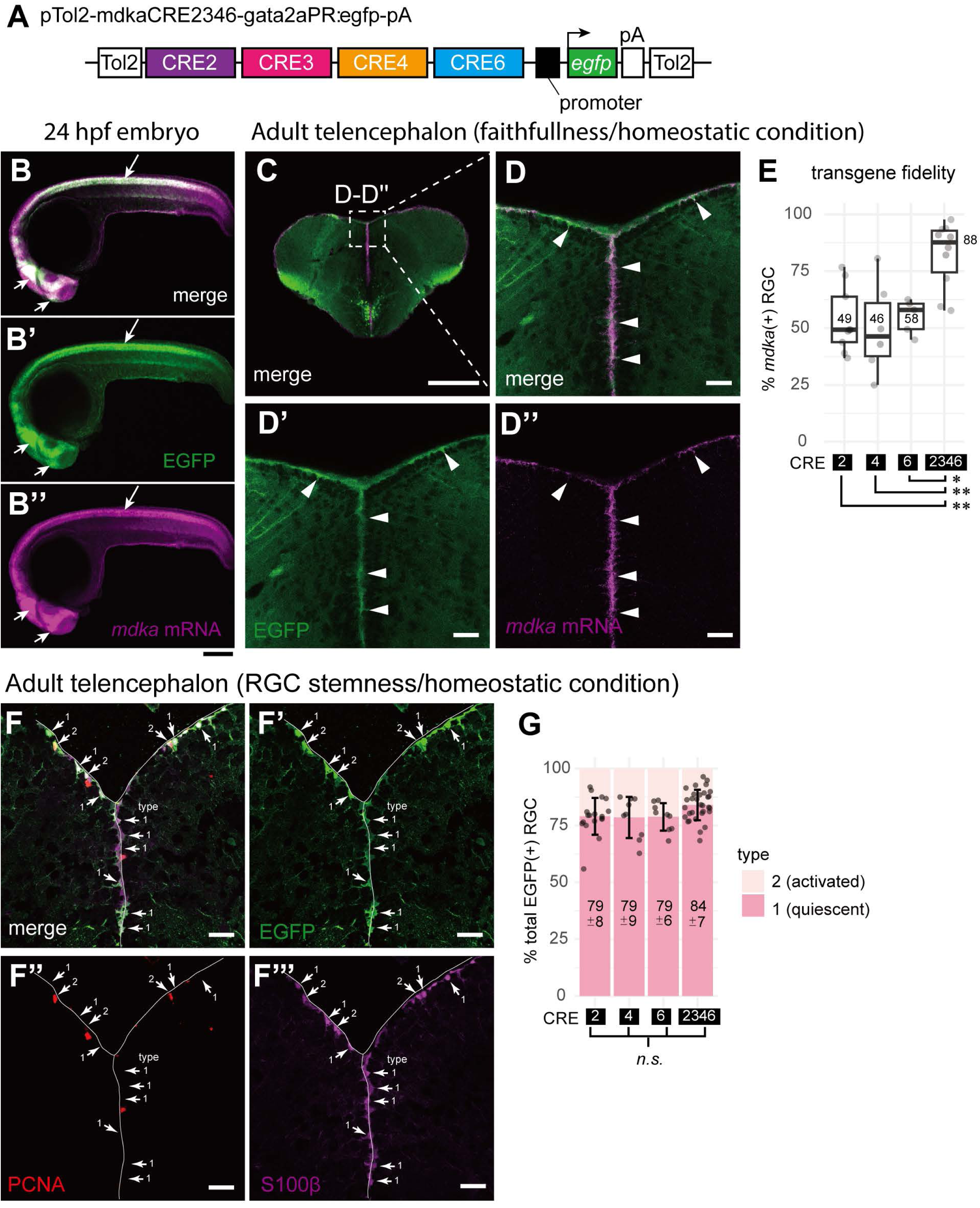
Combination of CREs enhances faithful transgene expression in both embryos and the adult telencephalon under homeostatic conditions. **A**, Schematic of the pTol2-mdkaCRE2346-gata2aPR:egfp-pA construct, in which a combination of CRE2, CRE3, CRE4, and CRE6 drives EGFP expression initiated by the *gata2a* promoter region (*gata2aPR*). **B-B’’**, Comparison of EGFP transgene (B’) and *mdka* mRNA (B’’) expression in 24 hpf embryo. Merged EGFP and *mdka* mRNA signals (B) show co-localization primarily in the brain and spinal cord (white arrows). **C**, Merged EGFP and *mdka* mRNA expression in a transverse section of the adult telencephalon under homeostatic conditions. The magnified Dm region (dashed rectangle) is shown in D-D’’. **D-D’’**, Magnified views of the telencephalic Dm domain showing merged signals (D), EGFP driven by CRE2346 (green, D’), and endogenous *mdka* mRNA (magenta, D’’). White arrowheads indicate RGCs in the ventricular zone (VZ). **E,** Faithfulness of EGFP transgene expression driven by CRE2, CRE4, CRE6 and the combined CRE2346. EGFP expression in VZ of each CRE transgenic line was compared with endogenous *mdka* mRNA. Data are shown as boxplots representing the percentage of EGFP⁺/*mdka*⁺ double-positive cells relative to total *mdka*⁺ cells. Median values are indicated. Significant differences were observed between CRE2346 and individual CREs (*ANOVA* followed by *Tukey’s* test; CRE2346 vs CRE2 [*p* < 0.01], vs CRE4 [*p* < 0.01**], vs CRE6 [*p* < 0.05*]). The data for individual CREs are identical to that shown in Figure 3K. **F-G**, Stemness analysis of RGCs labeled by CRE2346-driven EGFP. EGFP-positive cells in the VZ of the telencephalic Dm domain were categorized as type 1 (quiescent, S100β⁺/PCNA⁻) or type 2 (activated, S100β⁺/PCNA⁺) using immunolabeling. **F-F’’’**, Transverse telencephalon sections showing EGFP (F’), PCNA (F’’), S100β (F’’’) and merged image (F), highlighting type 1 and 2 RGCs under homeostatic conditions**. G,** Stemness of CRE2346-driven EGFP-positive RGCs was assessed and compared to those of individual CREs in a stacked bar chart. The individual CRE data are reused from Figure 3M. Mean ± standard deviation of type 1 RGCs (as a proportion of EGFP-positive RGCs) is shown. Each data point represents an individual brain slice. No significant differences were observed among different CREs (*n.s.*; *Kruskal-Wallis* test, *p* = 0.05). Scale bars: 200 µm (B-B’’ and C); 25 µm (D-D’’, F-F’’’). Sample sizes: *n* = 3 brains (C, D-D’’, E); *n* = 5 brains (F-F’’’, G).

We next investigated how the combination of *mdka* cis-regulatory elements (CREs) influences transgene expression during regenerative neurogenesis following brain injury (**Figure 6**). After stab injury, EGFP expression driven by the CRE2346 construct was observed in the ventricular zone (VZ) of both the injured (ipsilateral) and uninjured (contralateral) hemispheres, closely mirroring the endogenous *mdka* expression pattern (**Figure 6A, B-B’’**). Notably, injury-induced *mdka* upregulation at the lesion site, which occurs outside the VZ, was also faithfully reproduced by the CRE2346 line (**Figure 6C-C’’**). Quantification revealed that the number of radial glial cells (RGCs) expressing EGFP in the injured hemisphere was comparable to that in the contralateral side across all CRE lines, further reflecting endogenous *mdka* expression (**Figure 6D**). Moreover, EGFP intensity was significantly increased in the injured hemisphere in the CRE2346 line (*p* < 0.001), aligning with our prior observations of injury-induced *mdka* upregulation (**Figure 6E**) [21]. To evaluate the impact of combining CREs, we compared transgene fidelity and ectopic expression between CRE2346 and single CRE lines (**Figure 6F-G**). The combination significantly improved transgene fidelity, with consistent expression levels between ipsilateral and contralateral hemispheres (**Figure 6F**). Additionally, CRE2346 effectively suppressed the ectopic, off-target expression seen with CRE4 and CRE6 alone, which had previously shown high levels of non-specific activity (**Figure 6G**). We then assessed whether the CRE combination preserved the identity and stemness of transgene-expressing RGCs. Co-immunolabeling with S100β and PCNA allowed us to distinguish quiescent (type 1) from activated (type 2) RGCs (**Figure 6H-I’’’)**. In the uninjured (contralateral) hemisphere, CRE2346 increased the proportion of EGFP+ quiescent RGCs to 84%, closely approaching the 94% observed for endogenous *mdka* expression (**Figure 6J**). However, in the injured hemisphere, this fidelity was reduced. All constructs, including CRE2346, showed a marked decrease in the proportion of quiescent RGCs (39-69%), significantly lower than the 90% observed with endogenous *mdka*. This suggests that although the CRE2346 line closely replicates endogenous *mdka* activity under normal conditions and partially during regeneration, it likely lacks additional regulatory elements necessary for fully recapitulating *mdka*’s injury response, particularly for maintaining the quiescent RGC state.

**Figure 6.**
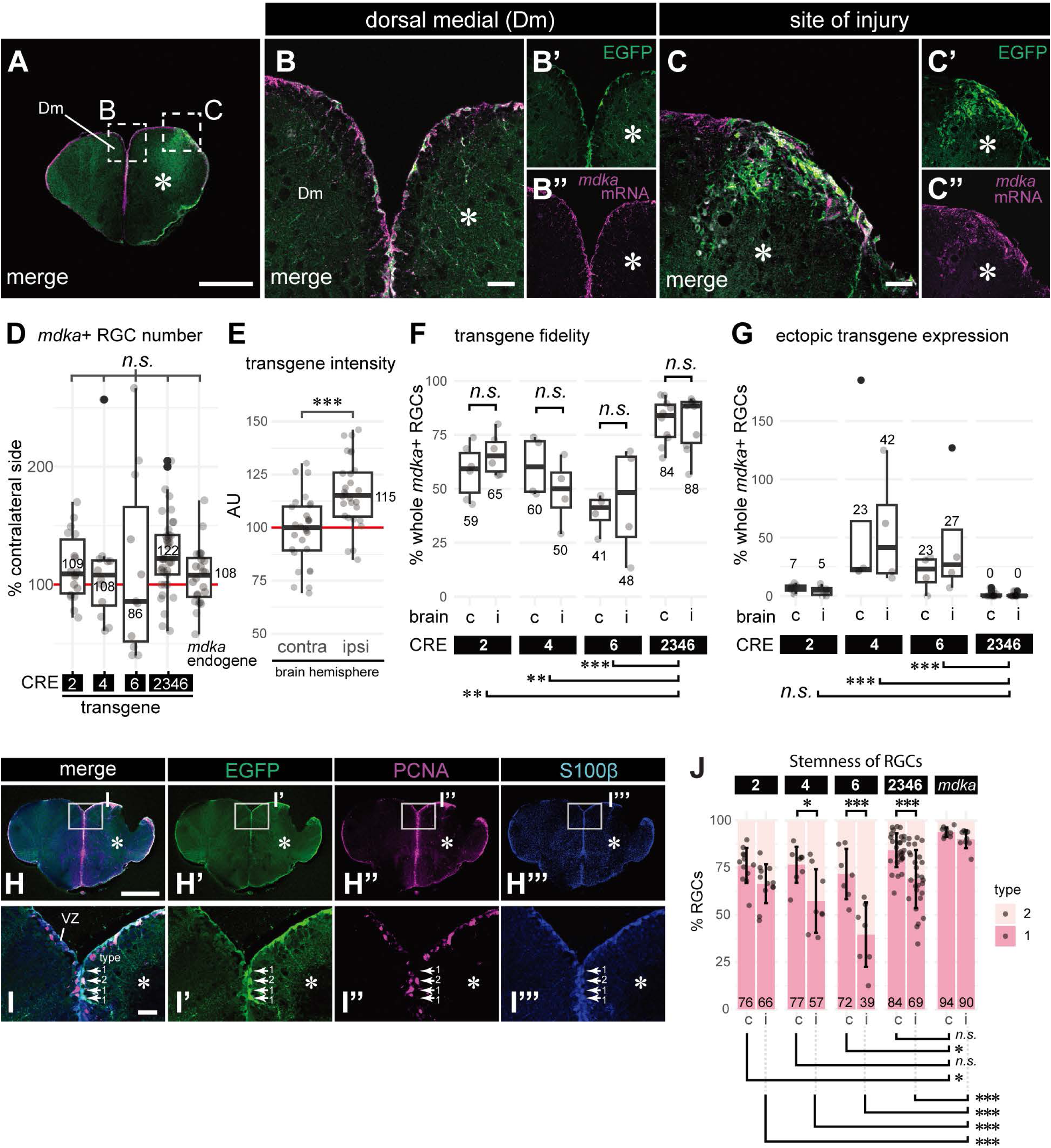
Combination of CREs enhanced faithful transgene expression in the adult telencephalon after brain injury. **A-C**’’Merged images of EGFP expression driven by CRE2346 (green) and endogenous *mdka* mRNA (magenta) in a transverse section of the adult telencephalon during regenerative neurogenesis at 5 dpl. The hemisphere ipsilateral to the injury is marked with an asterisk. **A**, Overview of the telencephalon. Stippled rectangular regions indicate areas magnified in B-B’’ (Dm domain) and C-C’’ (site of injury). **B-B’’**, Magnified merged image of CRE2346-driven EGFP (green, B’) and endogenous *mdka* mRNA (magenta, B’’). **C-C’**, Magnified merged view at the site of injury (C), showing upregulated expression of CRE2346-driven EGFP (green, C’) and endogenous *mdka* mRNA (magenta, C’’). **D,** The number of S100β-positive RGCs expressing either CRE transgenes or endogenous *mdka* mRNA in the hemisphere ipsilateral to the injury did not differ between CRE constructs and the endogenous gene (*n.s.*; *Kruskal-Wallis* test, *p* = 0.11). The number of ipsilateral RGCs is normalized to that of contralateral RGCs. **E**, In the CRE2346 transgenic line, EGFP levels in the ipsilateral hemisphere are significantly higher than in the contralateral side (*Welch t*-test, *p* < 0.001***). Normalized integrated fluorescence across brain slices is presented in arbitrary units (AU). **F-G**, Faithfulness of (F) and ectopic (G) EGFP transgene expression driven by either CRE2, CRE4, CRE6 or CRE2346 in contralateral (“c”) and ipsilateral hemispheres (“I”) after brain injury. **F**, EGFP expression in VZ of each CRE transgenic line were compared with endogenous *mdka* mRNA expression. Results are presented in boxplots showing the percentage of cells double-positive for EGFP and endogenous *mdka* mRNA relative to the total number of *Stemness*-positive cells. Significant differences were observed between CRE2346 and each individual CREs (*Dunn’s post hoc* test after *Kruskal– Wallis*: CRE2346 vs CRE2 [*p* < 0.01**], vs CRE4 [*p* < 0.01**] and vs CRE6 [*p* < 0.001***]). No significant difference was observed between contralateral and ipsilateral hemispheres in all cases (*n.s.*; *Dunn’s post hoc* test after *Kruskal–Wallis* analysis). **G**, Ectopic EGFP expression outside the VZ of each CRE transgenic line are shown relative to the number of endogenous *mdka*-positive RGCs. Although no significant difference was observed between contralateral and ipsilateral hemispheres, CRE2346 showed significantly lower ectopic transgene expression compared to CRE4 (Adjusted *p*-values in *Dunn* test following *Kruskal-Wallis* test; *p* < 0.001***) and CRE6 (*p* < 0.001***). **H-J**, Evaluation of RGC stemness in the CRE2346 line using PCNA and S100β immunostaining. **H-I’’’**, Transverse section of the adult telencephalon during regenerative neurogenesis at 5 dpl, showing the merged image (H-I) of CRE2346-driven EGFP (green, H’-I’), PCNA (magenta, H’’-I’’) and S100β (blue, H’’’-I’’’). The white rectangular regions in H-H’’’ are magnified in I-I’’’, respectively. Representative type 1 (S100β⁺/PCNA⁻) and type 2 (S100β⁺/PCNA⁺) RGCs are indicated (I-I’’’). **J**, Stemness of CRE2346-driven EGFP-positive RGCs compared to individual CRE lines and endogenous *mdka* mRNA (“*mdka*”). Pairwise comparisons were performed using *Dunn*’s test following a significant *Kruskal–Wallis* test (*n.s.* for *p* >= 0.05, * for *p* < 0.05, ** for *p* < 0.01 and *** for *p* < 0.001). The results for hemispheres contralateral (“C”) or ipsilateral (“I”) to the injury are plotted adjacent to each other. The data for CRE2, CRE4 and CRE6 are identical to Figure 4I. Each data point represents an individual brain slice. Black solid dots indicate outliers. Scale bars: 200 μm (A, H-H’’’); 25 μm (B–B’’, C-C’’, I–I’’’). Sample sizes: *n* = 4 brains (A, B-B’’, C-C’’); *n* = 6 brains (CRE2346; D, H-H’’’, I-I’’’, J); *n* = 3 brains (E, F, G); *n* = 3 brains (endogenous expression of *mdka*; D, J).

In summary, our findings demonstrate that the CRE2346 combination effectively drives *mdka*-like EGFP expression in RGCs with high spatial specificity and responsiveness to injury (**Table 3**). This highlights the necessity of multiple regulatory elements for precise spatial and temporal control of *mdka* expression under both physiological and regenerative conditions. Interestingly, all four CREs contain multiple transcription factor binding sites, including robust AP-1 binding sites within their evolutionarily conserved regions (**Figure S5**). AP-1 is a well-known regulator of gene expression during regeneration across various tissues and species, including the zebrafish heart, fin, and tail [35, 44–48], supporting the potential role of these CREs in the regenerative response to injury.

**Table 3.** Summary of EGFP expression patterns observed in distinct *mdka* CRE transgenic lines.

| CRE(s) | Activity summary |
| --- | --- |
| CRE2 | Adult regulatory sequence; active in physiological conditions; best matching *mdka* expression |
| CRE3 | Predominantly active during embryonic development; overlaps with *mdka* expression; shows ectopic expression in the adult telencephalon in Dp and DI. |
| CRE4 | Adult regulatory sequence; active in NSCs; does not show a significant response to injury |
| CRE6 | Adult regulatory sequence; active in NSCs with ectopic activity in neurons; no significant response to injury |
| CRE2346 | Recapitulates *mdka* expression; EGFP reporter overlaps with *mdka* mRNA in both embryo and adult; inducible upon injury |

Note: Dp, dorsal pallium; DI, lateral pallium; NSCs, neural stem cells.

## Discussion

Our study reveals that *mdka* expression in the zebrafish central nervous system is regulated by a complex and highly context-dependent network of CREs. These elements act in a modular, combinatorial, and sometimes redundant fashion, reflecting a multifaceted regulatory architecture that governs *mdka* transcription across different developmental stages and physiological states. Consistent with previous reports, *mdka* displays dynamic, tissue-specific expression patterns. For example, in the adult zebrafish telencephalon, *mdka* is expressed in RGCs under homeostatic conditions [21], while in the retina, it is absent in resting stem cells but becomes induced in proliferating Müller glia following injury [29, 49]. Interestingly, in the injured telencephalon, *mdka* expression remains restricted to quiescent RGCs, underscoring its tight, tissue-specific transcriptional control [21].

### Modular and context-specific regulation of *mdka*

Our transgenic analyses demonstrate that *mdka* expression is orchestrated by multiple CREs, each contributing distinct spatiotemporal aspects to its regulatory program. Some elements, such as CRE5 and CRE3, are active primarily during embryogenesis. For example, CRE3 drives strong expression in the embryonic brain and spinal cord but fails to recapitulate specific adult brain patterns, instead causing ectopic and nonspecific activation (**Table 3**). In contrast, elements like CRE2 are active mainly in the adult telencephalon, particularly in RGCs located in the Dm of the VZ. This modularity is not limited to the brain. In the heart, previous studies showed that a transgene containing *mdka* intronic sequences (*mdka*_e2, overlapping our CRE3, **Table S1**) was active in the developing epicardium but inactive after adult injury [35]. Meanwhile, a downstream element (*mdka*_e2, overlapping CRE6, **Table S1**) was inactive during development but became induced in a subset of adult epicardial cells following injury [35]. These examples reinforce the idea that distinct CREs mediate *mdka* expression in different developmental or physiological contexts.

### Combinatorial input cooperative and redundant CRE activity

While individual CREs contribute to specific features of *mdka* expression, none alone were sufficient to fully recapitulate endogenous expression, particularly in response to injury. For instance, CRE2, CRE4, and CRE6 each drove weak expression in adult telencephalic RGCs, but none showed injury responsiveness on their own. However, when combined in a multi-CREs construct, these same elements collectively drove expression that closely mimicked endogenous *mdka*, including robust injury-induced activation. This supports a model in which full *mdka* expression depends on the cooperative action of multiple CREs. Such combinatorial regulation may reflect the integration of distinct signaling inputs across separate regulatory elements. One possibility is that different signaling pathways act through specific CREs, with their additive or synergistic activity required to achieve the proper spatiotemporal expression profile. In this context, the effects of different CREs may be additive meaning that multiple enhancers with similar expression patterns together drive expression to levels that individual elements cannot reach on their own. Alternatively, or additionally, a single signaling pathway might act on multiple CREs in parallel, with full activation only occurring when all responsive elements are engaged. This combinatorial strategy may also contribute to enhanced specificity by limiting enhancer activity to overlapping spatial domains, thereby reducing ectopic expression. The presence of conserved AP-1 binding motifs within CRE2, CRE3, CRE4, and CRE6 (**Figure S5A-D**), coupled with the established role of the AP-1 complex in regeneration [34, 44, 47], supports both scenarios. Importantly, our observations also reveal redundancy among CREs. Although each element has specific functions, some show overlapping activity. For instance, despite weak individual responses to injury, CRE2, CRE4, and CRE6 all target the same VZ region. Furthermore, CRISPR/Cas9-mediated deletion of a regulatory element in the regenerating fin did not abolish *mdka* expression, and regeneration proceeded without detectable patterning defects [34]. This suggests that redundant or compensatory enhancer activity may preserve gene expression when individual elements are disrupted. Such redundancy adds robustness to the regulatory system, ensuring consistent *mdka* activation even in fluctuating or damaged environments. Together, our results highlight a regulatory logic in which enhancer modularity, cooperation, and redundancy work in concert to control *mdka* expression with high precision and resilience.

### Broader implications for gene regulation and regenerative biology

The modular and distributed regulatory architecture of *mdka* is consistent with classical models of enhancer function. The *even-skipped (eve)* gene in *Drosophila*, for example, is expressed in a seven-stripe pattern during embryogenesis, with each stripe controlled by a separate enhancer integrating local signals [50, 51]. Similarly, in mammals, the neural crest gene *sox10* is regulated by multiple enhancers with overlapping and redundant functions, enabling robust expression across tissues and stages [52, 53]. These cases, like *mdka*, illustrate how enhancer modularity enables nuanced, context-dependent expression and buffering against perturbation. Our *mdka* reporter lines provide a useful system for dissecting transcriptional programs that govern adult neurogenesis and injury-induced gene activation. These tools could be employed to identify upstream regulators of regenerative gene expression and screen for factors that modulate regenerative capacity *in vivo*.

### Regulatory complexity vs. simplicity: A comparative perspective

Intriguingly, the complexity of *mdka* regulation contrasts with the simpler enhancer architecture observed for other genes expressed in the same cellular context. For example, *id1*, which is co-expressed with *mdka* in quiescent RGCs under both physiological and regenerative conditions, appears to be controlled by a single enhancer responsive primarily to BMP signaling [24, 25]. Despite their overlapping expression domains, *id1* and *mdka* rely on very different regulatory strategies. This contrast raises important questions: Why do some genes require complex enhancer networks while others are regulated through simpler mechanisms? Comparative studies between genes like *mdka* and *id1* could illuminate how enhancer complexity evolves and functions in the context of regeneration and plasticity.

### Limitations and future directions in defining the *mdka* regulatory landscape

Our primary goal was to identify and functionally validate CREs that positively regulate *mdka* expression. Using *in vivo* transgenesis in zebrafish embryos and adults, we discovered several elements that drive EGFP expression in patterns consistent with endogenous *mdka* activity. These findings provide a foundation for understanding how *mdka* is regulated during development and regeneration. However, a key limitation of our approach is that it is optimized to detect enhancer activity and does not capture negative regulatory elements such as silencers. Traditional reporter assays are unable to distinguish between a lack of enhancer function and active repression. For instance, CRE1 did not drive detectable EGFP expression on its own, but whether this indicates silencer activity or simply absence of regulatory function remains unresolved. To address this, future studies should explore combinatorial testing, such as integrating CRE1 with active enhancers like CRE2346, to determine if it can restrict or suppress ectopic expression. Such designs could uncover context-dependent repression that would remain hidden when elements are tested in isolation. Our data also underscore the modular nature of the *mdka* regulatory architecture, which involves synergy, additivity, and redundancy among CREs. Additive effects, where multiple elements with overlapping activity are needed to achieve full expression, should be distinguished from redundancy, where one CRE can compensate for the absence of another. Both features likely contribute to the robustness and precision of *mdka* expression in both physiological and regenerative contexts. A fuller understanding of this regulatory logic will require experimental strategies that can reveal both positive and negative regulatory interactions.

Another limitation relates to the use of fluorescent reporter systems. The relative stability of EGFP protein may result in prolonged signal even after *mdka* mRNA expression has subsided, potentially contributing to discrepancies in spatial localization. To address this, we performed *in situ* hybridization for *egfp* mRNA, which confirmed both expected expression in the dorsomedial region and ectopic expression in the parenchyma (data not shown). These findings further support the idea that certain silencer elements may be absent from our current constructs. Given its lack of activity in isolation, CRE1 remains a strong candidate for a negative regulatory element, and its function in combination with other enhancers will be a priority for future studies. In addition, mRNA-level analysis across multiple transgenic lines will be essential for identifying those that most accurately recapitulate endogenous *mdka* expression in radial glial cells. Collectively, these insights highlight the importance of both enhancer and silencer elements in shaping the spatial and temporal dynamics of *mdka* regulation under homeostatic and regenerative conditions.

## Materials and methods Zebrafish husbandry

All zebrafish lines were maintained at the European Zebrafish Resource Center (EZRC). Experiments were performed on 6- to 12-month-old wild-type fish or transgenic reporter lines (**Table S2**). Animal husbandry was carried out in accordance with permit guidelines under §11 TSchG (Regierungspräsidium Karlsruhe, Aktenzeichen 35–9185.64/BH KIT). Fish were kept according to the European Society for Fish Models in Biology and Medicine (EuFishBioMed) guidelines at the Karlsruhe Institute of Technology (KIT) [54], and animal experimentation adhered to German animal protection standards, approved by the Government of Baden-Württemberg (Regierungspräsidium Karlsruhe, Reference Numbers: 35-9185.81/G-214/21 and 35-9185.81/G-215/21). Efforts were made to minimize animal distress and reduce their number.

### Plasmids construction

Plasmids for enhancer/promoter reporter assays were generated using Gateway cloning and TOPO/TA cloning strategies. The primers used for PCR amplification of putative CREs, and promoter are listed in **Table S3,** while the coordinates and sizes of the CREs and the promoter are provided in **Table S1**. Most plasmids (**Table S4**) were constructed using commonly available vectors, with detailed protocols for Gateway cloning available as described in Kwan et al., 2007 [55]. The pTol2-mdkaCRE2346-gata2aPR:egfp-pA plasmid was created by combining four *mdka* putative CREs (CRE2, CRE3, CRE4, and CRE6) into the pT2KHGPzGATA2C1 destination vector and was synthesized by GenScript Biotech Corporation.

### Microinjection to generate transient and stable transgenesis

One-cell stage embryos were injected with 1 μl of 20 ng/μl Tol2 transposase mRNA, 1 μl of 10 % phenol red, 50 ng/μl plasmid DNA, and nuclease-free water (final volume 10 μl) directly into the yolk through the chorion. After injection, embryos were incubated at 28 °C in E3 medium with methylene blue. At 24 hpf, embryos were sorted based on EGFP fluorescence. Positive F0 founders were identified by EGFP expression, raised to sexual maturity, and outcrossed with wildtype zebrafish. Stable EGFP expression in ∼20 % of F1 progeny confirmed successful stable transgenesis.

### Fluorescent *in situ* hybridization (FISH) of whole-mount embryos

Embryos were collected at the desired developmental stage, and chorions were removed manually or with pronase (1:100 w/v, 28.5 °C). The *mdka* probe (Molecular Instruments Inc.) was used, following the HCR™ RNA-FISH protocol with minor modifications, available on the Molecular Instruments website.

### Stab injury of adult telencephalon and preparation of sections

The stab injury experiment and telencephalon sections were performed following the protocol described by Schmidt et al. [43].

### FISH on adult telencephalon sections and embryos

For experiments using probes from Molecular Instruments Inc., the protocol “HCR™ RNA-FISH protocol for fresh frozen or fixed frozen tissue sections was followed, with minor modifications. Specifically, for 24 hpf embryos, proteinase K treatment was conducted at a concentration of 10 μg/ml (diluted from a 1000 × stock to 1 × working concentration) for 10 min to improve probe penetration. The *mdka* probe used for fluorescent *in situ* hybridization was purchased from the Molecular Instruments Inc. with a dilution of 1:500.

### Immunohistochemistry (IHC) on adult telencephalon sections

Vibratome sections were blocked with 1 ml blocking buffer for 1 hour at RT. After removing the buffer, sections were incubated overnight at 4 °C with primary antibodies in 300 μl blocking buffer. The next day, sections were washed 3 × 10 min with PTW (1 × PBS containing 0.1% Tween-20), then incubated with secondary antibodies (diluted in PTW) for 2 hours at RT in the dark. DAPI staining (1:2000 in PTW) was performed for 20 min if needed, followed by 3 × 5 min washes in PTW. Sections were mounted and imaged. All antibodies used are listed in **Table S5**.

### Mounting for imaging

For embryos, dechorionated embryos were embedded in a 1.6 % low melting-point agarose (Sigma) and 0.016 % Tricaine (Sigma) mixture in E3 medium, oriented within 1 min using a needle or fine hair, left at room temperature for 3 min, cooled at 4 °C for 3 min, supplemented with water drops to prevent evaporation. While for adult telencephalon sections, fluorescently labelled brain sections were mounted on glass slides with 0.17 mm coverslips using Aqua Polymount (PolySciences Inc., USA) and stored at 4 °C in the dark before and after imaging [43].

### Confocal imaging

Fluorescently labelled embryos and telencephalon sections were imaged using Leica TCS SP5 DM5000 and SP8/DLS confocal fluorescence microscope (Leica Microsystems, Wetzlar, Germany) equipped with 405 nm, 488 nm, 561 nm and 633 nm lasers. Excitation (Ex) wavelengths and emission (Em) ranges were as follows: Ex 488 nm for AlexaFluor 488 (EGFP, Em 492-550 nm), Ex 561 nm for AlexaFluor 561 (PCNA, Em 565-605 nm), Ex 633 nm for AlexaFluor 633 (S100β, Em 650-740 nm), Ex 633 nm for AlexaFluor 633-labeled *mdka* RNA probe (EM 655-724 nm), and Ex 405 nm for DAPI (Em 409-480 nm). Imaging was performed with sequential scanning at 16-bit color depth with a 1.0 AU pinhole aperture. Images were acquired at a resolution of 1024 × 1024-pixel resolution, with a pixel size of approximately 1.55 μm × 1.55 μm (field of view: ∼1.55 mm × 1.55 mm), using a 63 ×/1.20 NA Water Immersion objective (Leica Microsystems) at a scanning frequency of 400 Hz. Laser intensity, PMT gain, detector settings, AOTF (Acousto-Optic Tuneable Filters) attenuation and other imaging settings were kept constant across experimental groups to ensure comparability.

### Image processing and arrangement

Images were processed with consistent exposure times and gain settings across all series. Z-stack maximum projections were generated and cropping or rotation was performed in ImageJ/Fiji. For wide-field images in **Figure S2**, deconvolution using the Iterative Deconvolve 3D and Diffraction PSF 3D plugins were applied to remove out-of-focus blur and enhance clarity. All figures, including flowcharts, were created and arranged using Microsoft PowerPoint (version 16.78) or Adobe Illustrator 2022 (version 26.4.1).

### Statistical analysis

Three sections per brain were analyzed from at least three individuals (with sample sizes indicated in the figure legends). Each brain section was treated as a single data point. All statistical analyses were performed and visualized using R. To determine the appropriate statistical test, data normality was assessed using the *Shapiro–Wilk* test with a significance level of *α* = 0.05. For normally distributed data, parametric tests were used: the *Welch t*-test for comparisons between two samples, and the *Tukey HSD* test following a significant *ANOVA* (*α* = 0.05) for comparisons involving more than two groups. For non-normally distributed data, the *Mann–Whitney U* test was used for unpaired two-sample comparisons, and the *Dunn* test was applied following a significant *Kruskal–Wallis* rank sum test (*α* = 0.05) for multiple-group comparisons.

## Acknowledgements

We would like to sincerely thank Anne Schröck for her invaluable technical support throughout this project. We are grateful to Luisa Lübke, Gaoqun Zhang, and Barbara Schornick for their contributions during the early phase of this study, and to Fushun Chen for his initial support with bioinformatics. We also thank Ferenc Müller and Yavor Hadzhiev for their helpful discussions and advice on bioinformatics approaches. Research in Sepand Rastegar’s laboratory is supported by the BioInterfaces in Technology and Medicine and Natural, Artificial, and Cognitive Information Processing (NACIP) programmes of the Helmholtz Association, as well as by project grants from the German Research Foundation (DFG GRK2039, STR 439/17-1, RA 3469/5-1) and the Marie Skłodowska-Curie Initial Training Networks (ITNs) ZenCode and Danio-Recode. Jincan Chen is supported by a grant from the China Scholarship Council (CSC), grant number 202106310017. We thank the KIT Publication Fund of the Karlsruhe Institute of Technology for its support. Open access funding was enabled and organized by Projekt DEAL.

## Author contributions

SR designed, supervised the experiments, analysed the data, and co-wrote the manuscript with JC, MT and ND. JC, AH, MT and TB conducted experiments and analysed data. JC and MT analysed and quantified the data. CV conducted the Hi-C and 4C data analysis. All authors have reviewed and approved the final version of the manuscript for publication.

## Data availability statement

The data that supports the findings of this study are available in the main text and the supporting information of this article.

**Abbreviations**
CRE: cis-regulatory element
dpf: days post-fertilization
Dm: dorsomedial of the telencephalon
Dpl: days post-lesion
EGFP: enhanced green fluorescent protein
FISH: fluorescent *in situ* hybridization
hpf: hours post-fertilization
IHC: immunohistochemistry
kb: kilo bases
PCNA: proliferating cell nuclear antigen
RGCs: radial glial cells
mRNA: messenger
RNA: *mdka*: *midkine a*
NSCs: neural stem cells
S100β: S100 calcium-binding protein β
*Tg*: transgenic line
TSS: transcription start site
VZ: ventricular zone;

